# Effects of excess phosphate on a coastal plankton community: a mesocosm experiment in the Baltic Sea

**DOI:** 10.1101/2024.02.05.576994

**Authors:** Kristian Spilling, Mari Vanharanta, Mariano Santoro, Cristian Villena-Alemany, Matthias Labrenz, Hans-Peter Grossart, Kasia Piwosz

## Abstract

Eutrophication in the Baltic Sea has caused an imbalance in the inorganic nitrogen (N) to phosphorus (P) ratio, leaving excess phosphate (PO_4_) after the phytoplankton spring bloom that terminates after N-depletion. Using monitoring data, we demonstrated that the PO_4_ concentration has continued to increase in the outermost Gulf of Finland during past decades. We further investigated the fate of such excess PO_4_ in a two-week mesocosm (1.2 m^3^) experiment. The starting concentration of PO_4_ was 0.66 µM, and treatments included a non-treated control (control), nitrate addition (N-add; 3.6 µM), glucose addition (C-add; 25 µM) and combined nitrate and glucose addition (N+C-add). The addition of N both in N-add and N+C-add treatments stimulated nano- and microphytoplankton, while the picophytoplankton abundance increased only after N-depletion. Also, the copepod biomass was positively affected by the N-addition. N_2_-fixing cyanobacteria were present but in low abundance. Carbon addition did not enhance heterotrophic bacterial uptake of PO_4_ contrary to our expectations, nor did it affect the phyto- or zooplankton community composition. The PO_4_ concentration was reduced to ∼0.4 µM in the control and C-add treatments and to 0.16 µM in the two N-amended treatments, with an inorganic N:P uptake ratio of 6.7. These results underscore the role of picophytoplankton in reducing the excess PO_4_ pool after the spring bloom, a function traditionally ascribed to bloom-forming diazotrophic cyanobacteria in the Baltic Sea.

## Introduction

Deoxygenation, i.e. loss of oxygen, in the ocean is an increasing global problem that has direct implications not only for aerobic organisms but also for biogeochemical cycles (Keeling et al. 2010). There are two primary causes of the deoxygenation of the ocean: global warming, and human input of nutrients to marine ecosystems leading to eutrophication. Warming of surface waters happens on a global scale and reduces gas solubility, including oxygen. Eutrophication is a local or regional environmental problem that causes algal blooms that typically aggravate oxygen consumption in deeper water layers after the bloom, when the biomass is decomposing while sinking. In relatively shallow coastal seas, such elevated oxygen consumption may cause hypoxic and anoxic conditions on the seafloor, where sediment microbial processes typically decrease bioavailable nitrogen through denitrification and anammox (Hannig et al. 2007). At the same time, anoxia increases the release of phosphate from the sediment through the chemical reduction of iron-bound phosphate. The net result of anoxic bottoms is often a decrease in bioavailable nitrogen and an increase in phosphate concentration, causing the inorganic N:P ratio to decrease in several parts of the world (Conley et al. 2002, Hauss et al. 2012, Kalvelage et al. 2013, Meyer et al. 2016).

The Baltic Sea has been affected by eutrophication for several decades, which has increased the area of anoxic seafloor (Conley et al. 2002). In the Baltic Proper and the Gulf of Finland, the most productive period with accompanying flux of organic material to the sea floor is during the spring bloom, which in these subbasins terminate when dissolved inorganic nitrogen (DIN) has been depleted. There was a regime-like shift in the mid-1990s when the dissolved inorganic phosphorus (DIP) pools started to be at elevated levels after the spring bloom; low phosphate concentrations were generally detected during early summer before the mid-1990s, though N-limitation of the spring bloom could be already demonstrated at that time (Tamminen 1995). The decreasing inorganic N:P ratio is caused primarily by a release of phosphate from the seafloor after turning anoxic (Conley et al. 2002). At present the surplus of phosphate after the spring bloom typically range from 0.1 up to 0.6 µmol PO_4_ L^-1^ in the Gulf of Finland and the Baltic Proper (Spilling et al. 2018). However, there is relatively little data published in recent years, making it challenging to ascertain whether this trend persists.

The excess phosphate is believed to directly benefit N_2_-fixing cyanobacteria that often create toxic or nuisance blooms during the summer (Larsson et al. 2001, Kangro et al. 2007), supported by model work (Munkes et al 2021). *Nodularia spumigena* and *Aphanizomenon flos-aquae* are amongst the typical dominating filamentous cyanobacteria, forming blooms in July and August in large parts of the Baltic Sea. These species are N_2_-fixers and can take up PO_4_ and store it in the form of polyphosphate. However, they grow relatively slowly in water temperatures below 15°C (although *A. flos-aquae* is less sensitive to cold water than *N. spumigena*), and typically start forming blooms only after the excess phosphate pool has been depleted (Lehtimäki et al 1997, Wasmund 1997).

An alternative pathway of the excess P pool is going through other primary producers or heterotrophic bacteria (Nausch and Nausch 2004), thereby reducing the available P for the filamentous cyanobacteria. However, the prevailing N-limitation restricts the growth of other phytoplankton, whereas heterotrophic bacteria, that may also take up P, are typically limited by carbon during late spring (Lignell et al. 1992). Organic carbon produced by the spring bloom phytoplankton triggers a bacterial response, dependent on the composition of the phytoplankton community, but this labile carbon is rapidly consumed (Camarena-Gómez et al 2018, 2019). After this labile carbon is depleted, there remains a large (compared to other coastal ecosystems) refractory pool of dissolved organic carbon due to the high freshwater influence in the Baltic Sea bringing in e.g. humic substance (Hoikkala et al 2015).

Currently, it remains unclear which organisms take up the excess phosphate, most of which seems to settle out of the euphotic zone before cyanobacterial blooms develop (Nausch et al. 2008). Consequently, other P sources must exist for these cyanobacteria (Raateoja et al. 2011), most likely through upwelling, turbulent mixing events, or regeneration of organic P (Raateoja et al 2011, Wasmund et al. 2012).

Recently, we demonstrated in a small-scale (20 L) experiment that the post-spring-bloom plankton community could remove excess phosphate, even without any N-addition, but with a clear effect of temperature that affected both the rate of phosphate uptake and the plankton community composition (Vanharanta and Spilling 2023). Picophytoplankton seems to play a major part in the drawdown of phosphate, but an open question is the role of heterotrophic bacteria that can be competing with phytoplankton for this resource (Lignell et al. 1992, Vanharanta and Spilling 2023).

Here, we present an updated dataset of phytoplankton pigments and inorganic nutrients from the Finnish monitoring program taken in the Gulf of Finland, together with an experiment where we addressed the potential of an early-summer plankton community to remove the excess phosphate. The experiment was done in larger scale mesocosms closer to natural conditions than our previous indoor tank experiment (Vanharanta and Spilling 2023). In addition to a non-treated control, we added bioavailable nitrogen (N-add), carbon (C-add) and both (N+C-add), to stimulate primary producers and heterotrophic bacteria. We hypothesized that picophytoplankton would become relatively more important after N-depletion, taking up part of the excess phosphate, but that heterotrophic bacteria can also exploit the inorganic P pool when they have access to a labile carbon source.

## Materials and methods

### Field data

Field data was obtained from the Algaeline data constituting two commercial ships passing through the Gulf of Finland (Helsinki - Travemünde, and Helsinki – Stockholm, Supplementary Figure S1). The data consists of discrete samples automatically taken from a flow through system on the return to Helsinki where they were collected and analyzed on the same day. The variables measured include Chlorophyll a (Chla) concentration and inorganic nutrients. For Chla, GF/F filters (Whatman, UK) were used and the Chla was extracted in EtOH, stored at -20°C and in the darkness for up to two months, and measured with a spectrofluorometer (Varian Cary Eclipse, Agilent, USA), calibrated against known standards (Merck, Sigma-Aldrich), with excitation and emission wavelengths of 430 and 670 nm, respectively (Jespersen and Christoffersen 1987). Inorganic nutrients were determined using standard colorimetric methods (Grasshoff et al. 1999). The ship-of-opportunity monitoring data are available from 1998 until present. In addition, fluorescence sensor data in the flow-through system exists from 2013.

### Mesocosm experiment

The experiment was set up in 1.2 m^3^ mesocosm bags (Ø = 90 cm, length = 190 cm) that were moored outside the Tvärminne Zoological Station, Gulf of Finland (59° 50’ 40” N, 23° 14’ 57” E). The bags were attached to a floating platform (Spilling et al. 2022a). In total, 14 bags were installed but the two outermost bags were not used. These ‘dummy’ bags enabled to create the same light conditions for the outermost experimental units. All units were covered to prevent rain or bird droppings from falling into the bags. The spring bloom in the area had peaked in April and the PO_4_ concentration was relatively low at the onset of the experiment (0.013 µ mol L^-1^), thus we added PO_4_ to all units to a final concentration of 0.66 µ mol L^-1^. The experimental setup consisted of an unamended control, a treatment with added nitrate (N-add; 3.6 µmol N L^-1^), a treatment with added carbon in the form of glucose (C-add, 36 µmol C L^-1^), and a treatment with both nitrate and glucose addition (N+C-add). The rationale behind the treatments was to stimulate the phytoplankton with nitrate and heterotrophic bacteria with glucose. Each treatment had three replicates, and in total 12 bags were used for the experiment.

The bags were filled with surface water using gentle pumping on 7^th^ June 2021, which was considered day -1 of the experiment. This filling method had visually (microscope) been tested and found not to affect zooplankton but prevents small fish from entering the mesocosms. The following day (day 0), nutrients and carbon sources were added in the morning and the first full sampling took place right after the addition. The temperature was approximately 18°C and the salinity was 5.5 at the start of the experiment (Supplementary Figs S2 and S3). Water samples were taken daily with a Limnos water sampler (Hydro-Bios, Germany) from 0.5 m depth from the middle of the bags for fluorescence measurements and flow cytometer counts. CTD profiles were also taken daily from the bags and outside the experimental units, and we had dissolved oxygen loggers (HOBO U26; Onset Inc. USA) mounted inside each mesocosm bag. Rate of change in dissolved oxygen was calculated by linear regression during daytime (09 – 17) and nighttime (23 – 03). In addition to the daily sampling (e.g. CTD profile), larger volumes (5 L) were taken on regular sampling days: 0, 1, 3, 6, 8,10, 13 and 15. During these days, additional variables such as inorganic nutrients and particular organic carbon, nitrogen and phosphorus were measured.

Total Chla fluorescence was determined from each mesocosm bag by daily use of a handheld fluorometer (AquaPen, Photon Systems Instruments, Czech Republic), and direct Chla measurements were carried out by filtration in duplicates on regular sampling days, and analyzed as described above.

We used standard colorimetric methods (Grasshoff et al. 1999) using a photometric analyzer (Aquakem 250, Thermo Scientific, USA) to measure dissolved inorganic nutrients: nitrite + nitrate (NO_2_ + NO_3_), phosphate (PO_4_) and dissolved silicate (Dsi). Ammonium (NH_4_) was measured separately using a spectrophotometer (U-1100, Hitachi, Japan).

Particulate organic carbon (POC), nitrogen (PON) and phosphorus (POP) were determined from duplicate water samples filtered onto acid-washed (2M HCl for 15 min, then rinsed carefully with ultrapure water) and pre-combusted (450°C, 4 h) GF/F filters (Whatman, UK). POC and PON were measured with an element analyzer coupled with a mass spectrometer (Europa Scientific ANCA-MS 20-20 15N/13C, UK). POP was determined according to Solórzano and Sharp (1980) as modified by Koistinen et al. (2017). Samples for biogenic silicate (BSi) were filtered on polycarbonate membrane filters (0.8 μm, GVS, Italy) and measured according to Koistinen et al. (2018).

Abundance of nano- and microplankton (phytoplankton and microzooplankton) was determined with the FlowCam (Fluid Imaging, Yokogawa) from preserved (acid Lugol’s solution) samples. The samples were stored in a fridge and counted within a year from the sampling. We used the 10x magnification and 100 µm flowcell during the runs and the accompanying Visual Spreadsheet software for automatic calculation of biovolume. Further identification and visual inspection were done with an inverted microscope (Letiz Labovert) using a 40x objective.

Flow cytometer counts of live samples were done daily with a Sysmex (Japan), Partec – Cube 8 equipped with two lasers (488 and 561 nm), two scattering (forward and side) and three fluorescence detectors (610/30; 661/16; and 670/40, corresponding to the detection of phycoerythrin, phycocyanin, and Chla, respectively). The trigger was set on Chla fluorescence (670 nm). On each sampling day, we made several measurement-runs with beads and blanks to ensure correct particle counts. Size fractionations (0.8, 1, 2, 5, 10, and 20 µm filters) were also used to identify the approximate size of different phytoplankton groups. The scattering and fluorescence properties were used to gate different phytoplankton groups using the FCS Express 6 software. Four groups were identified: picoeucaryotes: size <2 µm with only Chla, Cryptophyte-like >2 µm with Chla and phycoerythrin fluorescence, nanophytoplankton size 2–20 µm and microphytoplankton: size >20µm with only Chla fluorescence.

*Synechococcus*-like cells, <2 µm with phycocyanin fluorescence, were counted at the same time as heterotrophic bacteria, using a LSR II, BD flow cytometer (Biosciences, USA) equipped with a 488 nm laser. These samples were fixed with 1 % paraformaldehyde (final concentration) for 15 min in darkness, flash frozen in liquid nitrogen, and stored at -80°C until analysis. Before measurements, samples were thawed and stained with SYBRGreen I (Molecular Probes, Eugene, OR, USA) at a 10^-4^ (v/v) concentration and incubated for 15 min in the darkness before counting. We used CountBright beads (Molecular Probes) in each sample to determine the measured volume. FACSDiva Software (BD Biosciences) and Flowing Software version number 2.5.1. were used to gate and obtain the cell counts.

Zooplankton was collected three times, at the start, middle and end of the experiment. Samples were taken with a plankton net (25 cm diameter, 50 µm mesh size) carefully dragged from the bottom of the mesocosm bags to the surface. Samples were immediately preserved with acid Lugol’s solution and stored at 4°C. Zooplankton was identified to genus level from images created using a flatbed scanner and analyzed at 20 x magnification.

### Data analysis

A one-way analysis of variance (ANOVA) was performed to test whether significant effects could be revealed by the treatments. Tukey’s Honest Significant Difference (HSD) *post hoc* test was performed to compare different treatments and assess the significance of differences between them after ensuring normality and homogeneity of the residuals. In case of the data following a non-normal distribution, the ANOVA was performed on ranks followed by Dunn’s *post hoc* test. All statistical tests were performed using SigmaPlot 15 software (Systat Software).

## Results

### Field data

From the Gulf of Finland monitoring data, the Chla fluorescence had a clear peak in spring at around April 20 (Day 110), followed by a clear water period after mid-May (Supplementary Fig S4). Phycocyanin fluorescence was more variable, elevated levels of phycocyanin fluorescence started to appear after mid-May (around May 19, Day 140) but the regression we used indicated a peak around mid-July (July 14, Day 195), when the average temperature reached 15°C.

From the discrete samples, the Chla peak during spring was around 20 µg Chla L^-1^, with some observations of even higher concentrations, up to 33 µg Chla L^-1^ (Supplementary Fig S5). Nitrate was rapidly depleted during spring, with very low concentrations after the Chla peak. Phosphate was also reduced, but often not completely depleted, and there was more phosphate in the water during summertime (Day 138-220) in the 10-year period 2014-2023 compared with the other periods 1998-2003 and 2004-2013 (Dunn; p <0.002). There was no difference in phosphate concentration between the two earliest periods (Dunn; p >0.9). The concentration of dissolved silicate decreased during spring but never completely depleted, and the concentration was higher in the past decade (2014 - 2023) compared to the previous one (2004 – 2013; Dunn p <0.001).

### Mesocosm experiment

In the experiment, there was a clear effect of N addition, but no effect of carbon addition, on the phosphate concentration (Fig 1). The nitrate added to the N-add and N+C-add treatments was depleted already on day 3. The phosphate concentration decreased from ∼0.66 µmol L^-1^ at the start of the experiment to ∼0.16 µmol L^-1^ in the two treatments with N added on Day 8, resulting in an inorganic N:P uptake ratio of 6.7. In the control and C-add treatments, the PO_4_ concentration was 0.33 and 0.36 µmol PO_4_ L^-1^, respectively, at the end of the experiment. There was residual ammonium with a concentration in the range of 0.15 to 0.35 µmol L^-1^ throughout the entire experiment. Dissolved silicate was constant in the control and C-add but decreased slightly towards the end of the experiment in the N-add and N+C-add treatments, with an uptake of approximately 1.9 µmol L^-1^ (Fig 1).

**Fig 1.**
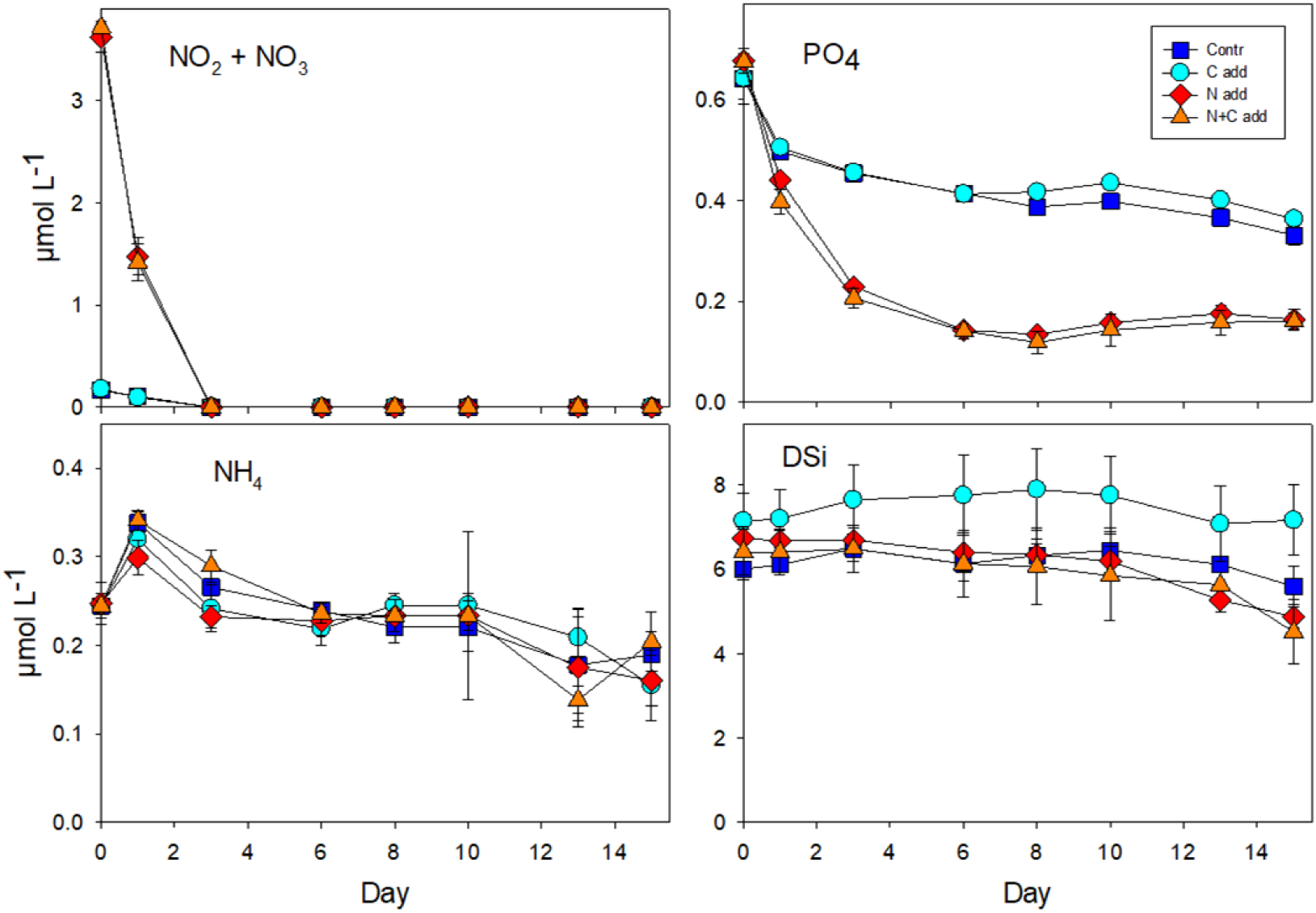
The concentration of nitrite + nitrate (NO_2_ + NO_3_), phosphate (PO4), ammonium (NH_4_) and dissolved silicate (DSi) plotted against time. The error bares represent the standard error (SE, n = 3).

The phytoplankton biomass declined from the onset of the experiment in the control and C-add treatments but with a delay of two days in the two N amended treatments (Fig 1). The initial Chla concentration was 3.5 µg L^-1^ and it decreased to 1–2 µg Chla L^-1^ and within two days in the control and C-add treatments (Fig 2). After reaching a minimum of <1 µg L^-1^, the Chla concentration increased again at the end of the experiment to ∼2 µg Chla L^-1^.

**Fig 2.**
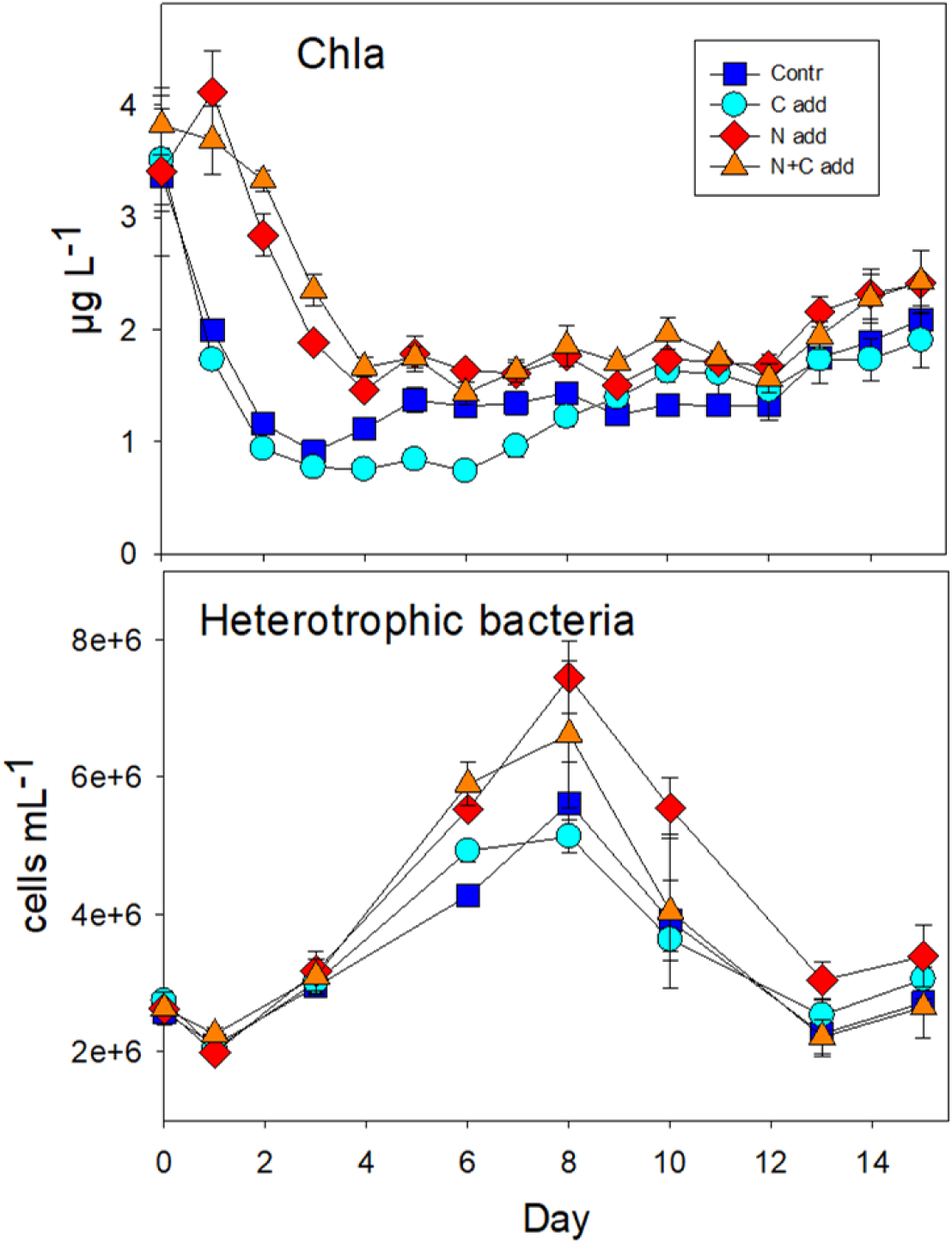
The concentration of Chlorophyll a (Chla) and abundance of heterotrophic bacteria plotted against time. The error bares represent the standard error (SE, n = 3).

Oxygen dynamics indicated more O_2_ was added during daytime in the N-add treatment compared with the N+C add treatment during the first three days (Supplementary Fig S6). During nighttime, the loss of O_2_ was higher in the N-add and N+C add treatments compared to the control and C-add treatments at the start of the experiment, but O_2_ dynamic was similar in all treatments after day 7 (Supplementary Fig S6).

Heterotrophic bacteria showed the same dynamic in all treatments (Fig 2); their abundance increased after day 1, with a peak at day 8. The overall bacterial abundance was higher in the N-add treatment compared to the control and C-add treatments (Tukey: p = 0.001 and p = 0.003 respectively), the rest of the treatments were similar (Tukey; p > 0.14).

All particulate nutrients decreased in the experimental units during the first week, but the decrease was more rapid in treatments without any nitrate addition (control and C-add), except for biogenic silicate (BSi) where the decrease was similar in all treatments (Fig 3). Stoichiometric ratios were relatively stable with an average C:N:P ratio of 140:14:1, and without any differences between treatments (ANOVA; p >0.1).

**Fig 3.**
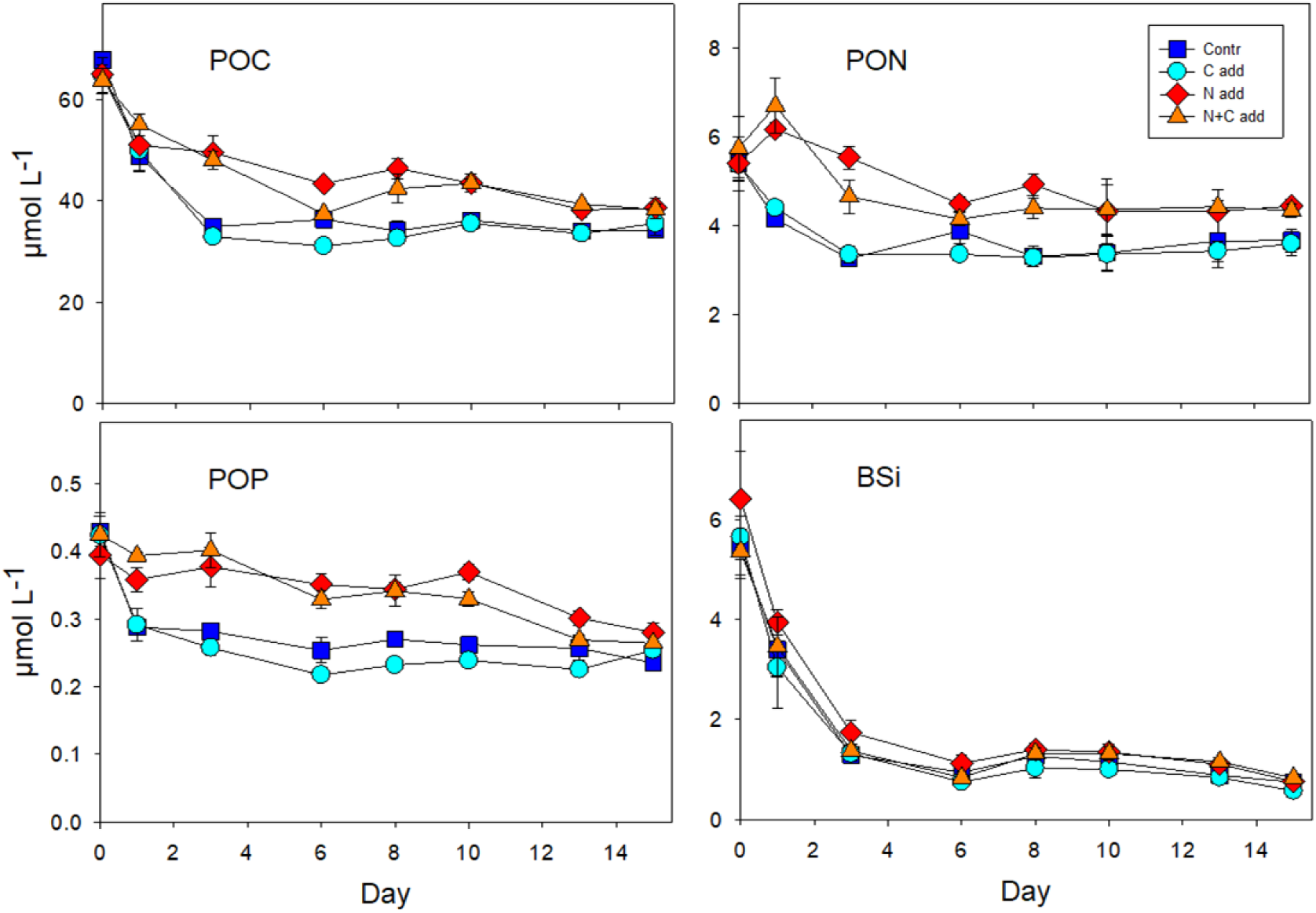
The concentration of particulate organic carbon (POC), nitrogen (PON), phosphate (POP), and biogenic silicate (BSi) plotted against time. The error bares represent the standard error (SE, n = 3).

The temperature and salinity throughout the experiment are presented in Supplementary Figs S2 and S3. There was a drop in temperature after day 6 from ∼18°C to ∼14°C due to upwelling, followed by increase towards the end of the experiment to ∼20°C. Salinity varied outside the bags but stayed constant within the bags with no sign of mixing with the outside water indicating that our bags remained intact during the experiment.

### Plankton community composition

Nitrate additions benefited micro- and nanophytoplankton that increased in abundance in both the N-add and N+C-add treatments (Fig 4), contributing to a high Chla concentration over the first three days. This was also evident from the FlowCam images, where, *e.g.,* the cell concentration and total biovolume of dinoflagellates >10 µm increased in these two treatments until nitrate had been depleted (supplementary Fig S7). The green algae *Monoraphidium* sp. (nanophytoplankton) increased also in the N-amended treatments, but kept growing after N depletion, reaching a maximum biovolume at day 8 (Fig 5).

**Fig 4.**
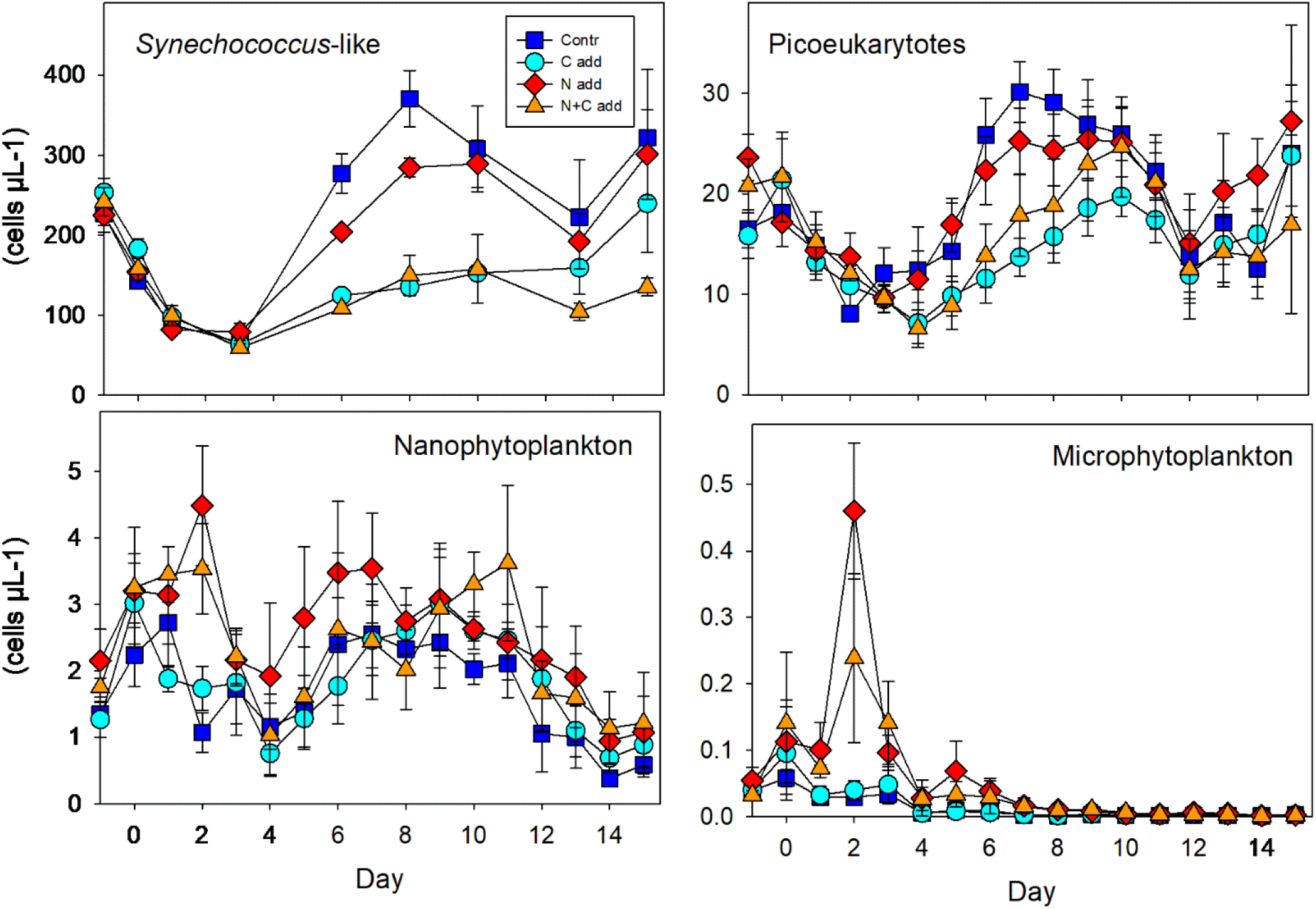
The abundance of *Synechoccoccus*-like cells, picophyto(?)eukaryotes, nano- and microphytoplankton plotted against time. The error bares represent the standard error (SE, n = 3).

**Fig 5.**
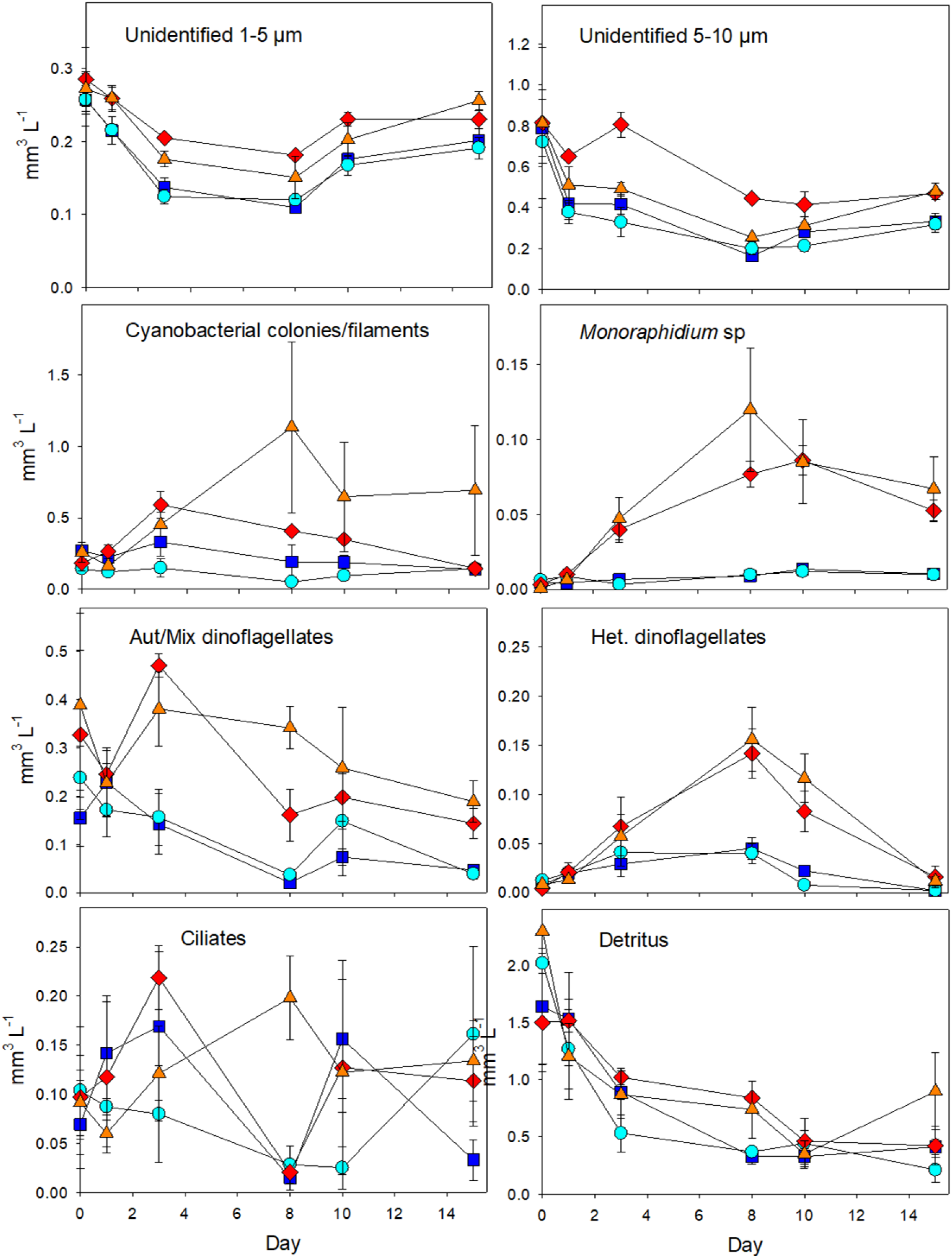
The biovolume of different nano- and microplankton groups plotted against time. The error bars represent the standard error (SE, n = 3).

After nitrate depletion, picoeukaryotes started to increase in abundance, especially in the control and N-add treatment, but also, to a lesser extent, in the N+C-add and C-add treatments (Fig 4). *Synechococcus*-like cells decreased in all treatments after the start of the experiment but increased again after day 3 and more so in the control and N-add treatments (Fig 4), peaking on day 8 when the abundance was clearly higher in the control and N-add treatments compared with the carbon amended treatments (Tukey p <0.014). *Planktothrix* sp., and *Pseudoanabaena* sp. were the main component of colonial and filamentous cyanobacteria. They peaked in the N-add and N+C-add treatments on day 3 and 6, respectively, with the highest peak in biovolume in the N+C-add treatment (Fig 5). *Aphanizomenon flos-aquae* and *Nodularia spumigena* were present but contributed combined less than 5% to the total cyanobacterial biovolume.

Microzooplankton consisted of heterotrophic dinoflagellates and ciliates (Fig 5). Heterotrophic dinoflagellates increased in the N-add and N+C-add treatments with a peak on day 8 before declining to a similar biovolume as in the control and the C-add treatments at the end of the experiment (Fig 5). The biovolume of ciliates was highly variable between replicates, and although the average biovolume was higher in the two N-add treatments, there was no consistent difference between the treatments (ANOVA, p = 0.767).

The mesozooplankton community was dominated by copepods, cladocerans and pelagic stages of *Amphibalanus* sp. (Fig 6). The number of copepods was highest in both N-add and N+C-add treatments towards the end of the experiment including both copepodite and nauplii stages (ANOVA, p = 0.024 and p = 0.017, respectively). Among cladocerans, the abundance of *Bosmina* sp. was positively affected by the nitrate addition (ANOVA, p = 0.04), whereas *Podon* sp. (Fig 6) and *Evadne* sp. (data not shown) both decreased over time and had completely disappeared in all treatments at the end of the experiment. The difference in *Amphibalanus* nauplii abundance between treatments was not clear (ANOVA, p = 0.06), but cypris larvae had only developed in the two N-amended treatments at the end of the experiment (Fig 6).

**Fig 6.**
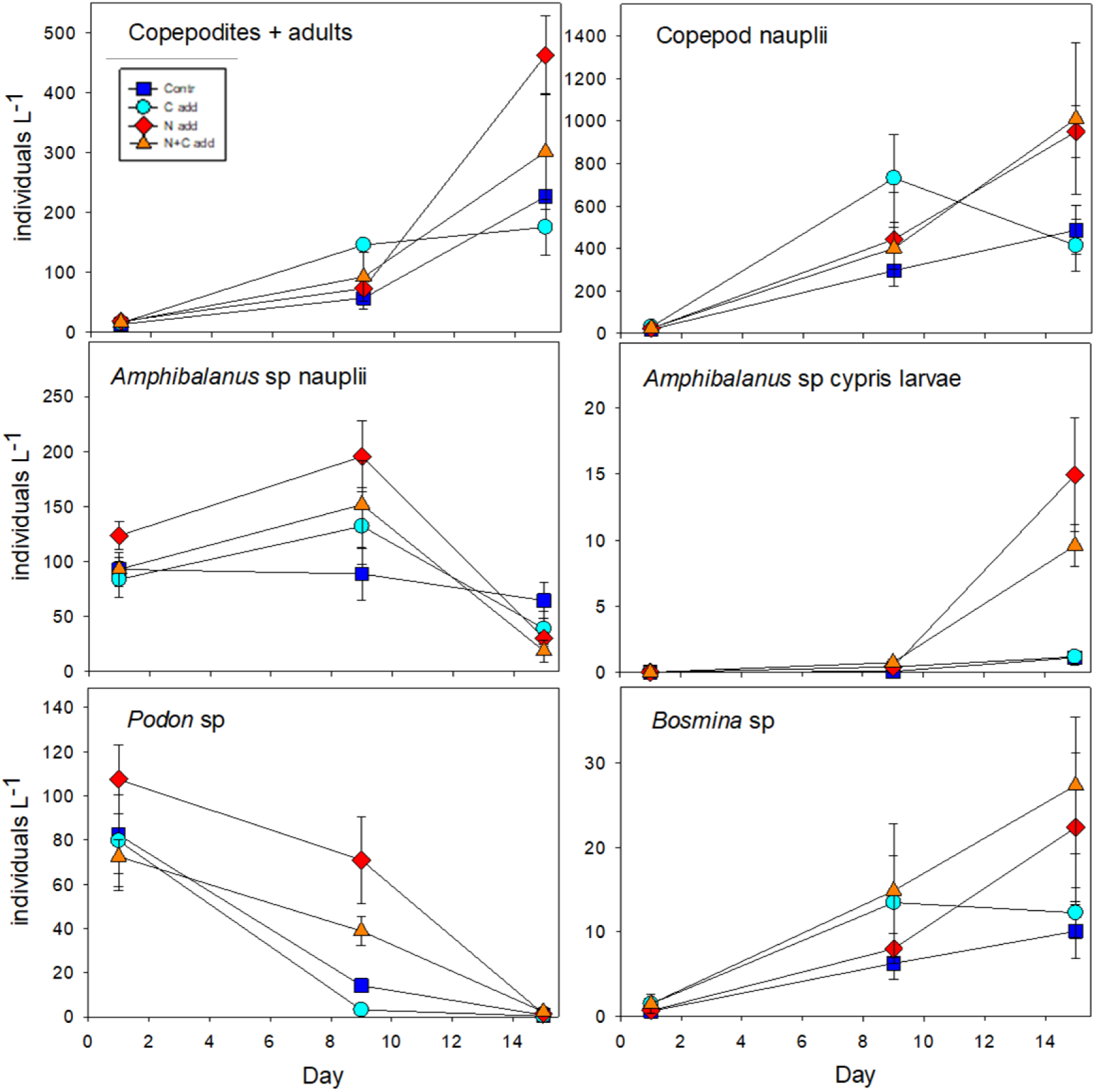
The abundance of different mesozooplankton groups plotted against time. The error bars represent the standard error (SE, n = 3).

## Discussion

Here we demonstrate that the phosphate concentration was higher in the past decade compared to earlier periods in the Gulf of Finland, suggesting that, despite a considerable reduction in the external nutrient load over past decades, it is still increasing. The reason for the high phosphate concentration is the release from the sediment (Pitkänen et al 2001). The effect of short-term shifts in the inorganic N:P ratio depends to some extent on the plankton community composition, but tends to be reflected in the organic N:P ratio (Spilling et al. 2019, Bach et al. 2020). Yet, potential long-term consequences of this phenomenon are still not known. A decreasing inorganic N:P ratio during N-depletion, as we found here, is often caused by reduced O_2_ concentrations in different ecosystems. However, it is not a universal process, as there are examples of other processes that increase N:P ratio, for example, by more effective P-removal from wastewater, increased use of N-rich fertilizer, or increased atmospheric N-deposition (McQuatters-Gollop et al. 2007, Kim et al. 2014). Despite all the efforts to reduce external loading to the Baltic Sea, the increasing excess phosphate concentration after N-depletion demonstrates that improving ecological state takes a long time to achieve, and arguably longer than it takes to deteriorate the ecosystem.

We also found that dissolved silicate had increased during the active growth season for algae. This is somewhat surprising as the DSi concentration has decreased long term in the Baltic Sea, linked to a higher retention of silicate by damming of rivers and increased burial of biogenic silicate in the sediment due to eutrophication (Conley et al 2008). Increases in dissolved silicate could therefore be interpreted as a positive sign of slightly improving conditions (Wasmund et al 2017), but further studies are needed to investigate whether this is a general trend in the Baltic Sea.

### Temporal plankton community development in the experiment

Inorganic nutrients initially stimulated nano- and microphytoplankton, in particular the chlorophyte *Monoraphidium* sp. and autotrophic dinoflagellates. This was expected, as they have a larger volume for internal storage, i.e. luxury uptake of nutrients. In addition, *Monoraphidium* sp. is known to have a relatively high growth rate (Seip and Reynolds 1995). Nonetheless, this nutrient effect only delayed the gradual decline of phytoplankton biomass. After inorganic nutrients had been depleted, there was a clear increase in picophytoplankton abundance, which has been observed in previous mesocosm experiments in this area (Crawfurd et al 2017, Spilling et al. 2022b). Smaller cells have a larger surface-to-volume area and are better adapted to low inorganic nutrient concentrations, which is the likely reason for this group to dominate during periods when easily available nutrient sources have been depleted by larger cells.

The mesocosm bags prevent effective wind-driven mixing, although there was some temperature driven advection. There was particulate organic matter sedimentation during the experiment, but unfortunately, we could not sample the bottom part of the mesocosms to quantify it. This export loss might have had some consequences for the phytoplankton community composition as non-motile groups, such as diatoms, likely sedimented out quicker than motile flagellates. This could be deduced from the development in BSi over time, and diatoms were a minor group in terms of biovolume.

As expected, mesozooplankton required a longer time to respond to the experiment, yet, an effect of the nutrient addition on its biomass was measurable (Fig. 6). The bags were not very deep (1.9 m) and the available organic material in the bottom likely supported the community of mesozooplankton that could easily move vertically between different water depths, especially since higher predators were absent. This is a likely reason for a treatment effect on copepods and *Bosmina* sp, even though the available, suspended food items were similar in all bags after the drawdown of nitrate. Some groups, such as the cladoceran *Evadne* sp. disappeared over time, which could be due to an enclosure effect as this cladoceran often tend to disappear in mesocosm experiments (e.g. Graneli and Turner 2002).

### Treatment effects

The main driver for the observed differences between treatments was the addition of nitrate that directly affected the phytoplankton biomass. The carbon addition, on the other hand, had no effect on particular organic nutrients, nor on bacterial abundance, suggesting that the total amount of added labile carbon was relatively low compared to the extra phytoplankton-derived carbon in the N-amended treatments. Interestingly, however, carbon addition did have a negative effect on picoeukaryotes and *Synechococcus*-like cells relative to the other treatments. This could have been due to competition for nutrients with heterotrophic bacteria, which likely got a boost due to the sudden, easily available carbon source that was added. This was, however, not evidenced by the abundance of heterotrophic bacteria that remained similar between control and C-add treatments. Increased grazing pressure could have played a role, but no noticeable differences in protozoan or microzooplankton abundance were observed and grazing would be expected to affect autotrophic and heterotrophic picoplankton equally (Kuosa and Marcussen 1988). Thus, the underlying reason for the apparent carbon effect on picophytoplankton still remains unclear.

Carbon addition also affected the O_2_ dynamics during the first three days of the experiment. There was not as much increase in O_2_ during the daytime in the N+C compared to the N-add treatments, which likely was due to higher primary production in the latter treatment. The N amended treatments revealed an increase in biomass and O_2_ reduction rate during nighttime suggesting a higher O_2_ loss due to respiration than in the control and C-add treatments. Any treatment effects on O_2_ dynamics disappeared after the first week of incubation, indicating similar biological activity between all treatments during the final week of the experiment. The rate of change in O_2_ is not directly transferable to biological activity, however, as temperature fluctuations affect gas solubility, and there were occasions of both O_2_ out-gassing and in-gassing due to changes in temperature.

### Phosphate removal

In the N-add and N+C-add treatments, ∼75 % of the phosphate was removed, whereas in the control and C-add treatments ∼ 40 % was removed within a week. Approximately 15 % of the phosphate pool, or 20–30 % of the PO_4_ taken up was released into the dissolved organic phosphorus (DOP) pool, but DOP concentration decreased again during the last week of the experiment (Vanharanta et al 2024), suggesting the added DOP was relatively labile and microbially available. The N:P uptake ratio of 6.7, less than half the Redfield ratio of 16, suggests an active luxury uptake of P of the plankton community. The decrease in the phosphate pool was not seen as an increase in the particulate phosphorus fraction although the N-amended treatments had a higher concentration of particulate organic elements. This was likely due to the export of particulate organic matter to the bottom of the mesocosm bags.

In a recent study, using smaller 20 L tanks, we found that the excess phosphate had been depleted completely under N-deficiency, even without N addition (Vanharanta and Spilling 2023). This previous experiment had a longer duration (35 d) than the one presented here, with a drawdown of phosphate in the control units of 20 % to 50 % (from a starting concentration of 0.55 µM) after 15 days, depending on the temperature (Vanharanta and Spilling 2023). Thus, it is likely that we would have also seen a complete drawdown of phosphate, if the current experiment would have lasted a few weeks more. In addition, the decrease of temperature by several degrees in the middle of the experiment likely decelerated microbial activities and consequently phosphate uptake rates. Possibly, more of the excess phosphate would have been taken up if temperature had stayed stable during the whole duration of the experiment.

The addition of glucose did stimulate bacterial productivity and extracellular enzyme activity (Vanharanta et al. 2024a), but there was no indication of any additional drawdown of phosphate by heterotrophic bacteria when stimulated by glucose compared to the control. Also, in this respect, the combined N+C addition behaved like the N addition. This further supports that most phosphate uptake was due to picophytoplankton with little contribution of heterotrophic bacteria. The extracellular enzymatic activity of free-living bacteria likely served to acquire carbon from polymers released by phytoplankton which was demonstrated by higher cell-specific activities of proteolytic and glycolytic enzymes during the bacterioplankton peak in the N-addition when compared to other treatments (Vanharanta et al. 2024a). This finding demonstrates the dependency of heterotrophic bacteria on phytoplankton and could explain why carbon addition alone did not result in significantly higher phosphate uptake compared to the control. The timing of the heterotrophic bacterial abundance peak is typical following the spring bloom when there is lots of organic matter available (Camarena-Gómez at al. 2018), and the bacterial community is to some extent affected by the phytoplankton community during the spring bloom (Camarena-Gómez et al. 2021).

### Link to cyanobacterial blooms in the Baltic Sea

One of the most prominent effects of eutrophication in the Baltic Sea is the intensifying occurrence of diazotrophic cyanobacterial blooms during summer. The prevailing N-limitation leads to a low inorganic N:P ratio, which favors N_2_-fixing cyanobacteria during warm, calm summer months (Niemi 1979, Wasmund et al. 2005). These cyanobacterial blooms amplify the eutrophication problem, as the amount of cyanobacterial fixed N is comparable to the anthropogenic N-loading in parts of the Baltic Sea (Savchuk 2005, Wasmund et al. 2005). The cyanobacterial input of N might thus create a self-reinforcing cycle through a positive feedback loop, i.e. N_2_-fixation increases the biomass load reaching the sediment, which in turn raises oxygen consumption and releases additional P into the ecosystem (Vahtera et al. 2007a, Spilling et al 2018).

In our experiment we categorized the different cyanobacteria present in three functional groups. The first group consists of small, single-celled cyanobacteria, here termed *Synechococcus*-like cells, which do not fix nitrogen and are presumably functionally closer to picoeukaryotes than to the larger N_2_-fixing filamentous species. The second group includes the relatively small filamentous cyanobacteria *Planktothrix* sp. and *Limnothrix* sp. plus the larger *Pseudoanabaena* sp. Common for all of them is that they do not fix N_2_. The third functional group comprises the relatively large N_2_-fixing filamentous cyanobacteria *Aphanizomenon flos-aquae* and *Nodularia spumigena*. A surplus of P would benefit the latter, especially if they have been able to increase their biomass during the nitrate depletion phase. Although these cyanobacteria were present in all treatments, there was no indication that these typical bloom-forming diazotrophic species proliferated (staying below 5% of the cyanobacterial biomass). Rather, the additional P uptake after N depletion seems to have been driven by an increase in picoeukaryotes, *Synechococcus*-like cells and possibly the smaller (non-N_2_-fixing) filamentous cyanobacteria. These organisms can take up phosphate and store excess phosphorus as polyphosphate (Jentsch et al. 2023). This was also observed under the microscope, although we lack quantitative data on polyphosphate storage. *Planktothrix* sp., and *Pseudoanabaena* sp. species are common in the area during summer, but do not form extensive blooms.

*Aphanizomenon flos-aquae* and *Nodularia spumigena* have different P uptake strategies. *A. flos-aquae* can take up and store excess P for later growth and also take up organic forms of phosphorus, whereas *N. spumigena* more commonly takes up phosphate and, to some extent, relies more on recycled P after phosphate depletion (Hagström et al. 2001, Vahtera et al. 2007, Schoffelen et al 2022). Although both species were present in the experiment, they were at exceptionally low concentrations. Other filamentous cyanobacteria (*Planktothrix* sp., and *Pseudoanabaena* sp.) made up the bulk of the measured cyanobacterial biovolume. The lack of a typical cyanobacterial community composition could be due to an enclosure effect, but both. *A. flos-aquae* and *N. spumigena* have relatively low growth rates (Reynolds 1984, Vahtera et al. 2005), and it is likely that they did not grow fast enough in the experiment to produce much biomass in the relatively short time span of the experiment. This is similar to the typical natural development where the excess phosphate typically has been depleted before the onset of filamentous cyanobacterial blooms later in summer. Our experimental results support our hypothesis that picophytoplankton plays a key role in the uptake of the excessive phosphate after the spring phytoplankton bloom. This would indicate that in contrast to current models/assumptions, the excess phosphate remaining after the spring bloom do not directly benefit the typical bloom forming cyanobacteria (*A. flos-aquae* and *N. spumigena*), which would rather rely on recycled or upwelled sources of phosphate.

In conclusion, we found that N-addition stimulated both nano- and microphytoplankton, and the picophytoplankton abundance increased following nitrate depletion. Based on these results, we conclude that picophytoplankton incorporate substantial parts of the excess P, which strengthen our initial hypothesis. Yet, picophytoplankton was unable to completely deplete the inorganic P-pool during the relatively short experiment. Most of the organic P formed was likely exported by the overall decrease in total phosphorus combined with a negligible change in the DOP pool between the start and the end of the experiment. Carbon addition, in contrast, stimulated heterotrophic bacteria production but did not increase bacterial uptake of excess phosphate, contrary to our expectations. The typical bloom-forming cyanobacteria in the Baltic Sea, considered to greatly benefit from excess phosphate, were present but not abundant in our mesocosms. Our results underscore the key role of picophytoplankton in reducing the excess phosphate pool after the spring bloom, a function traditionally ascribed to bloom-forming diazotrophic cyanobacteria in the Baltic Sea.

## Acknowledgements

We would like to thank the staff at Tvärminne Zoological Station for all their support, technical assistance, and nutrient measurements. In particular Joanna Norkko, Jaana Koistinen, Mervi Sjöblom, Kia Rautava and field technicians Göran Lundberg and Jostein Solbakken. We would also like to thank Sebastian Ehrhart for help with the monitoring data.

This study used equipment part of the Finnish marine research infrastructure (FINMARI) consortium.

This work was supported by the Transnational Access program of the EU H2020-INFRAIA project (No. 731065) AQUACOSM - Network of Leading European AQUAtic MesoCOSM Facilities Connecting Mountains to Oceans from the Arctic to the Mediterranean - funded by the European Commission. Additional funding came from Walter and Andree de Nottbeck foundation (KS and MV) and the Research Council of Finland (KS, decision no 354272). KP was supported by the National Science Centre, Poland under the Weave program (project no 2021/03/Y/NZ8/00076). HPG was funded by the German Science Foundation (DFG) project PycnoTrap (GR1540/37-1). MS was supported by Leibniz Science Campus Phosphorus Research Rostock in the funding line strategic networks of the Leibniz Association.

## Data availability

Data from all measured variables during the mesocosm experiment are available on PANGAEA (Vanharanta et al. 2024b).

---

**Supplementary Fig S1.**
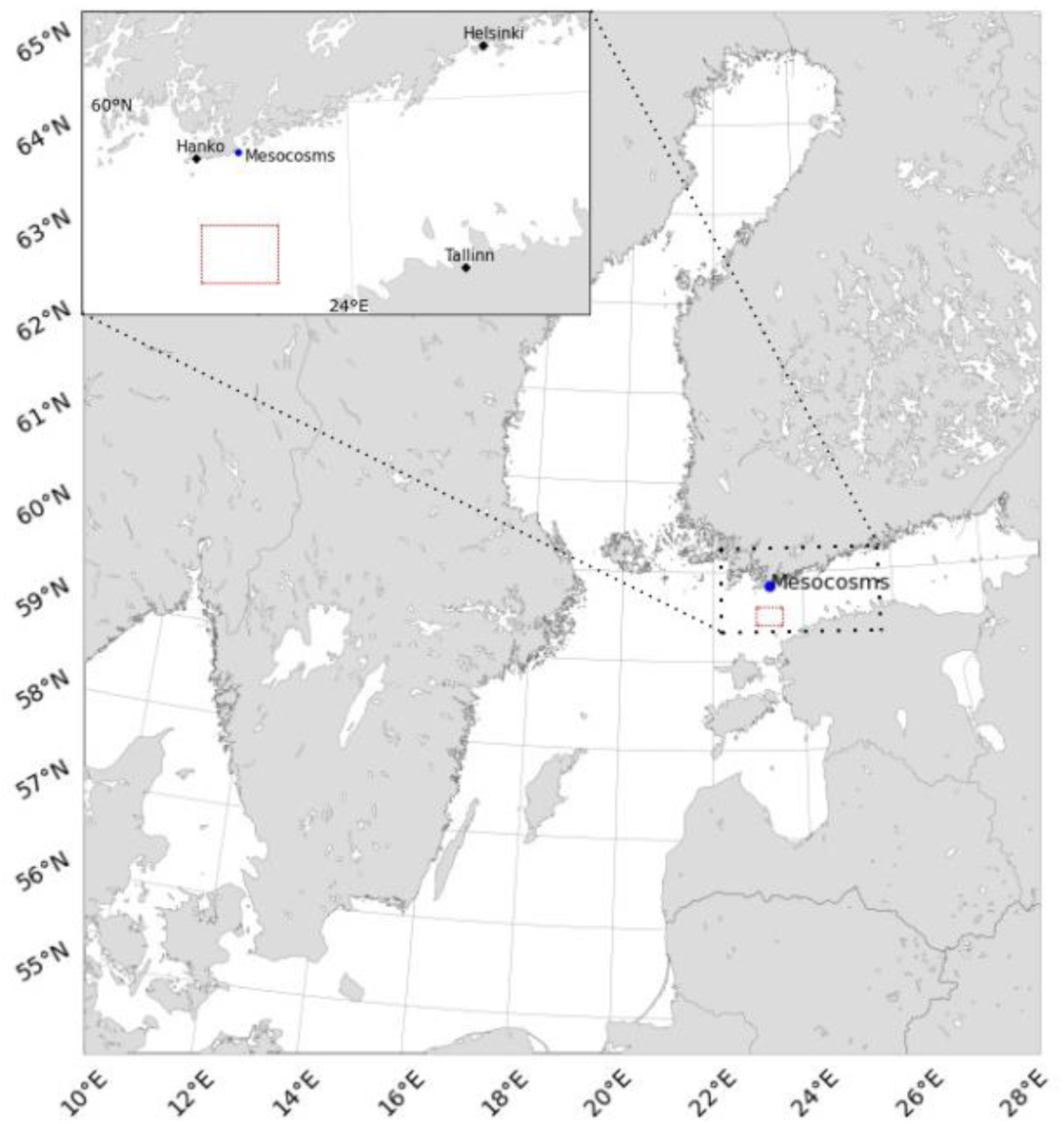
Map of the Gulf of Finland with an outline of where the field data was gathered from ships-of-opportunity (red box), and the mesocosm experiment (blue dot).

**Supplementary Fig S2.**
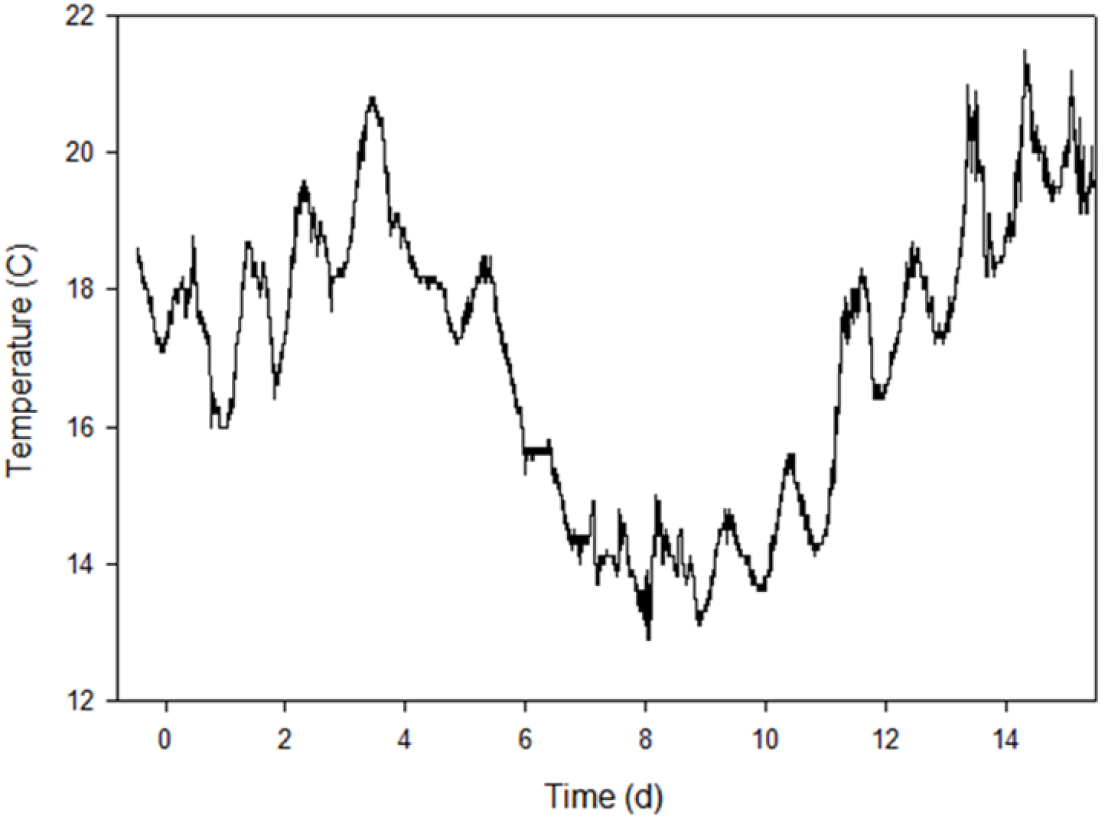
Temperature during the experimental setup. Temperature was measured every 30 s with a temperature probe placed at 50 cm depth right outside the mesocosm bags.

**Supplementary Fig S3.**
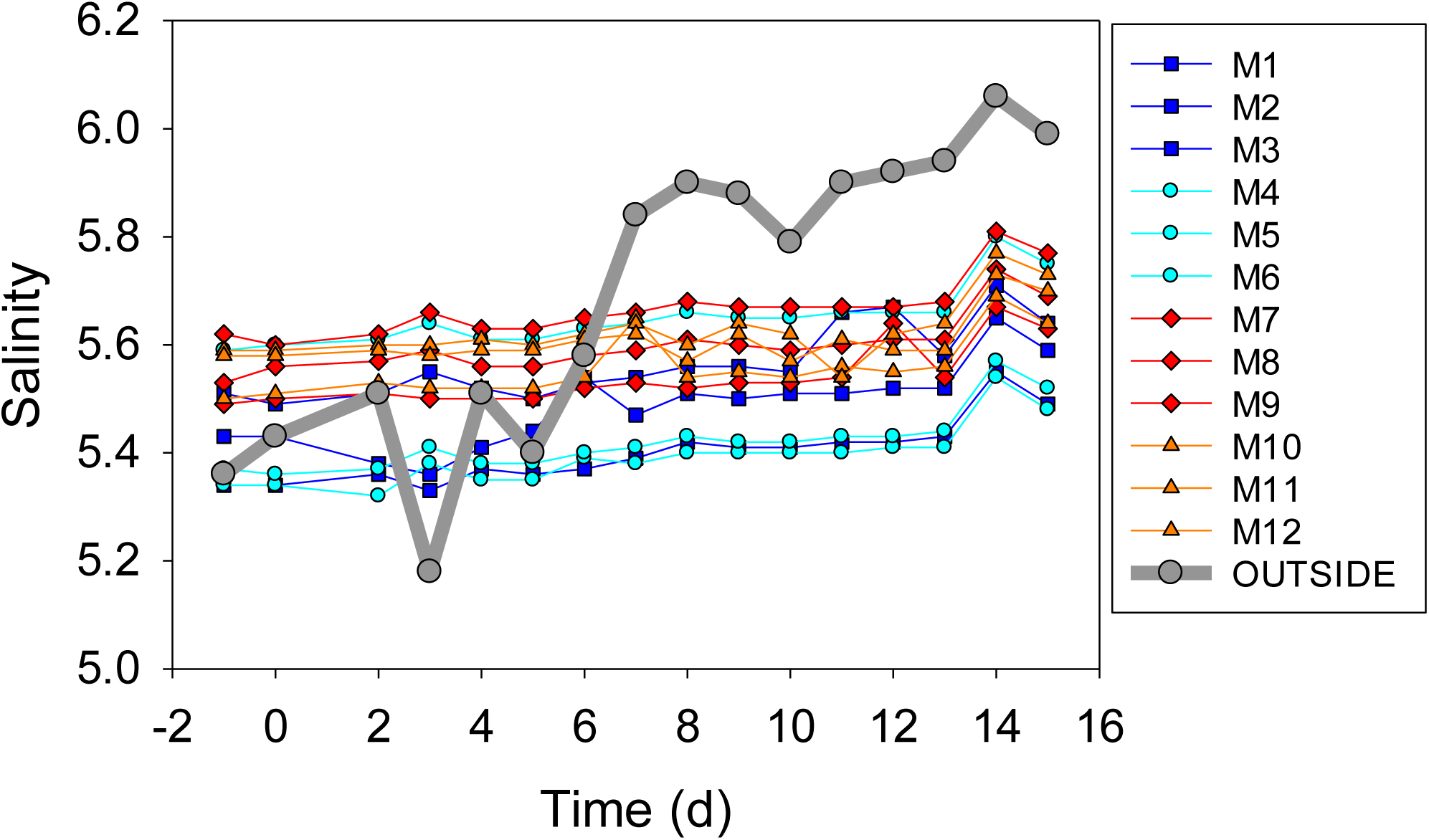
Salinity measured at 50 cm depth inside the 12 experimental bags, plus outside the mesocosm bags (thick grey line).

**Supplementary Fig S4.**
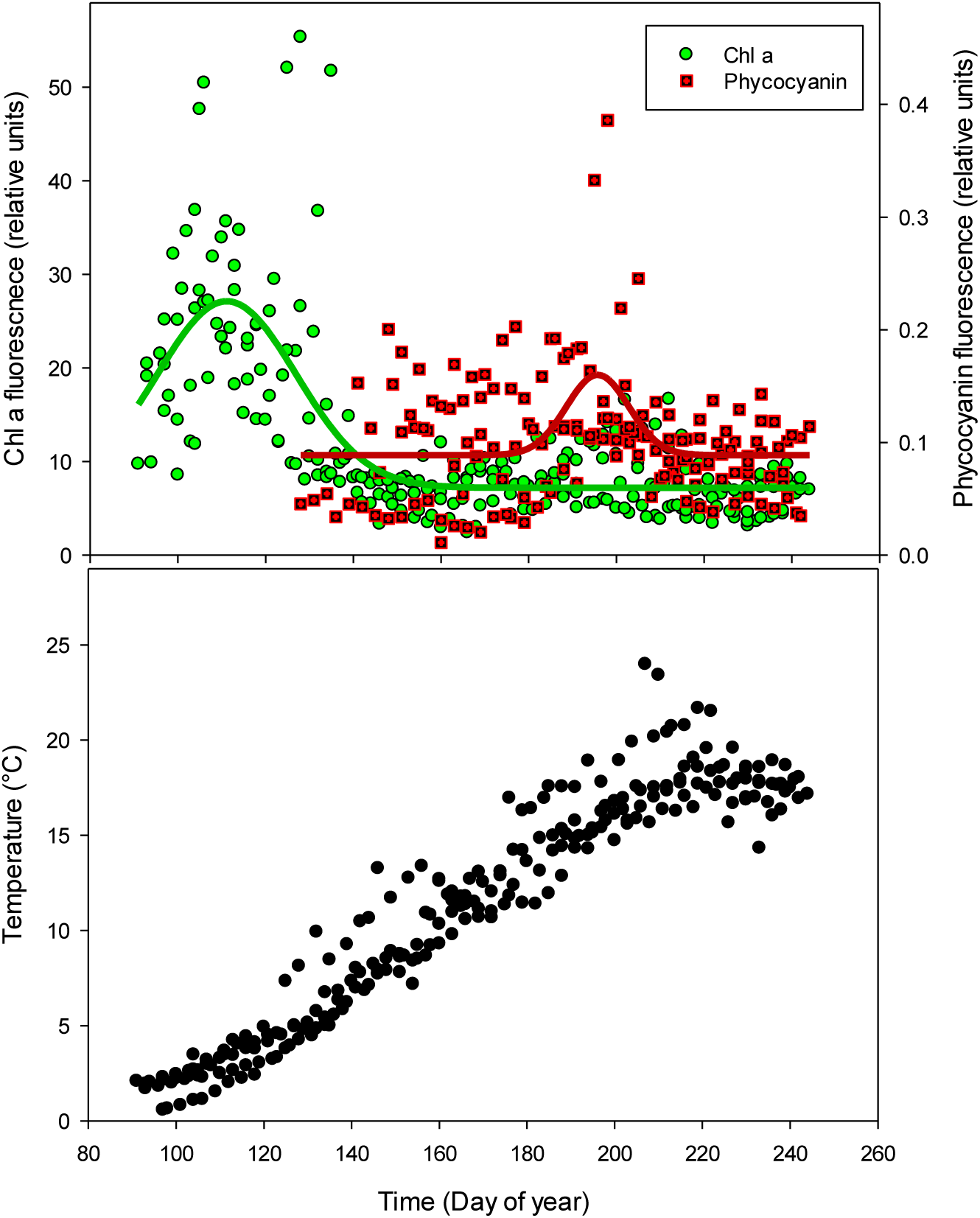
Chla and phycocyanin fluorescence (upper graph) and temperature (lower graph) taken outside the Hanko peninsula Gulf of Finland, from a flow through system on two different ships of opportunity. Each point represents an average from the area of one passing. Data from 2013 to 2016. Solid lines represent the Gaussian regression line.

**Supplementary Fig S5.**
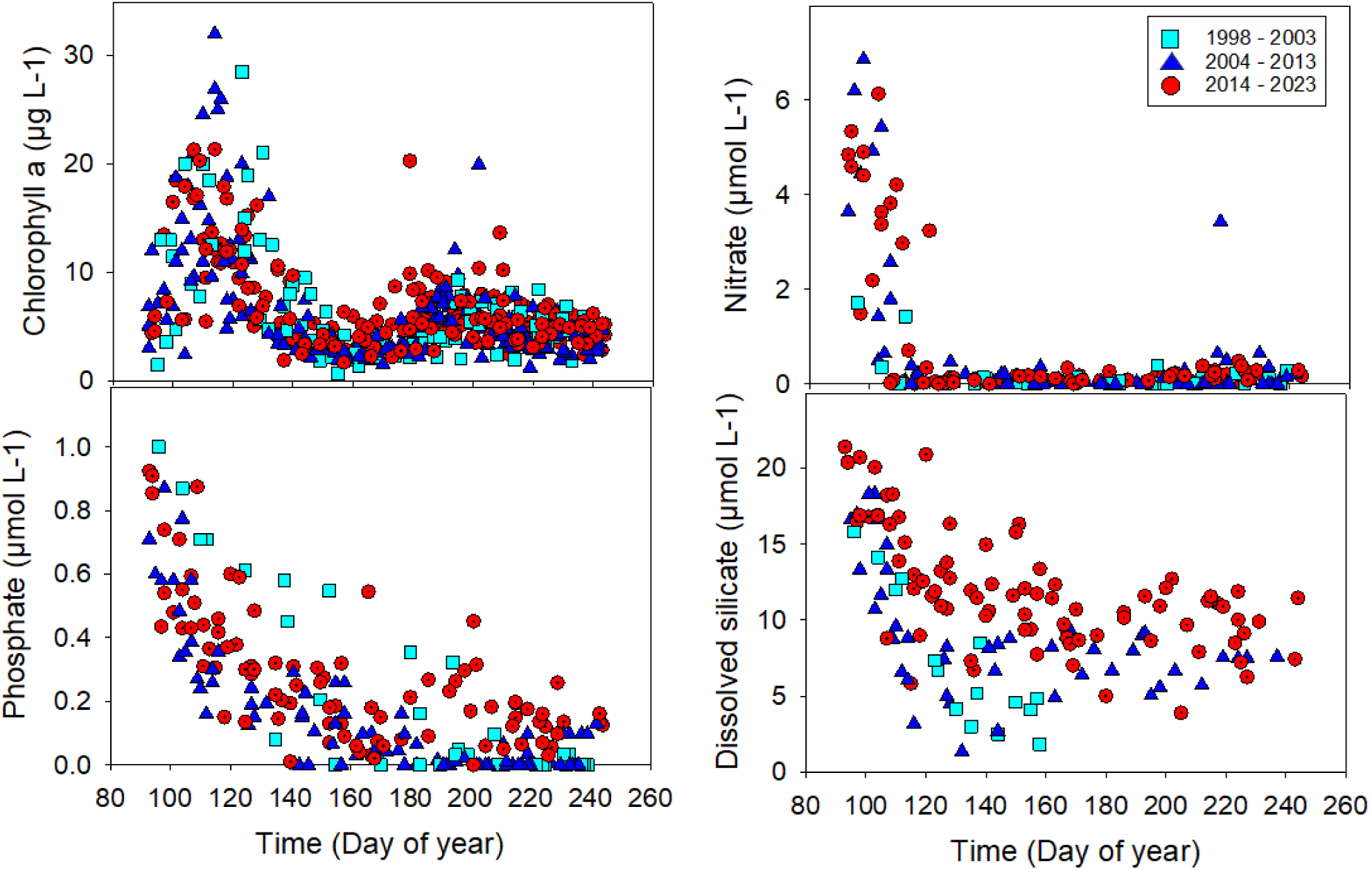
Chlorophyll and inorganic nutrients (nitrate, phosphate and dissolved silicate) taken from discrete sample taken outside the Hanko peninsula Gulf of Finland, in a flow through system on two different ships of opportunity. Each point represents one bottle filled from the transect. Colors represent three different periods, cyan square = 1998 – 2003, blue triangle = 2004 – 2013 and red circle 2014 – 2023.

**Supplementary Fig S6.**
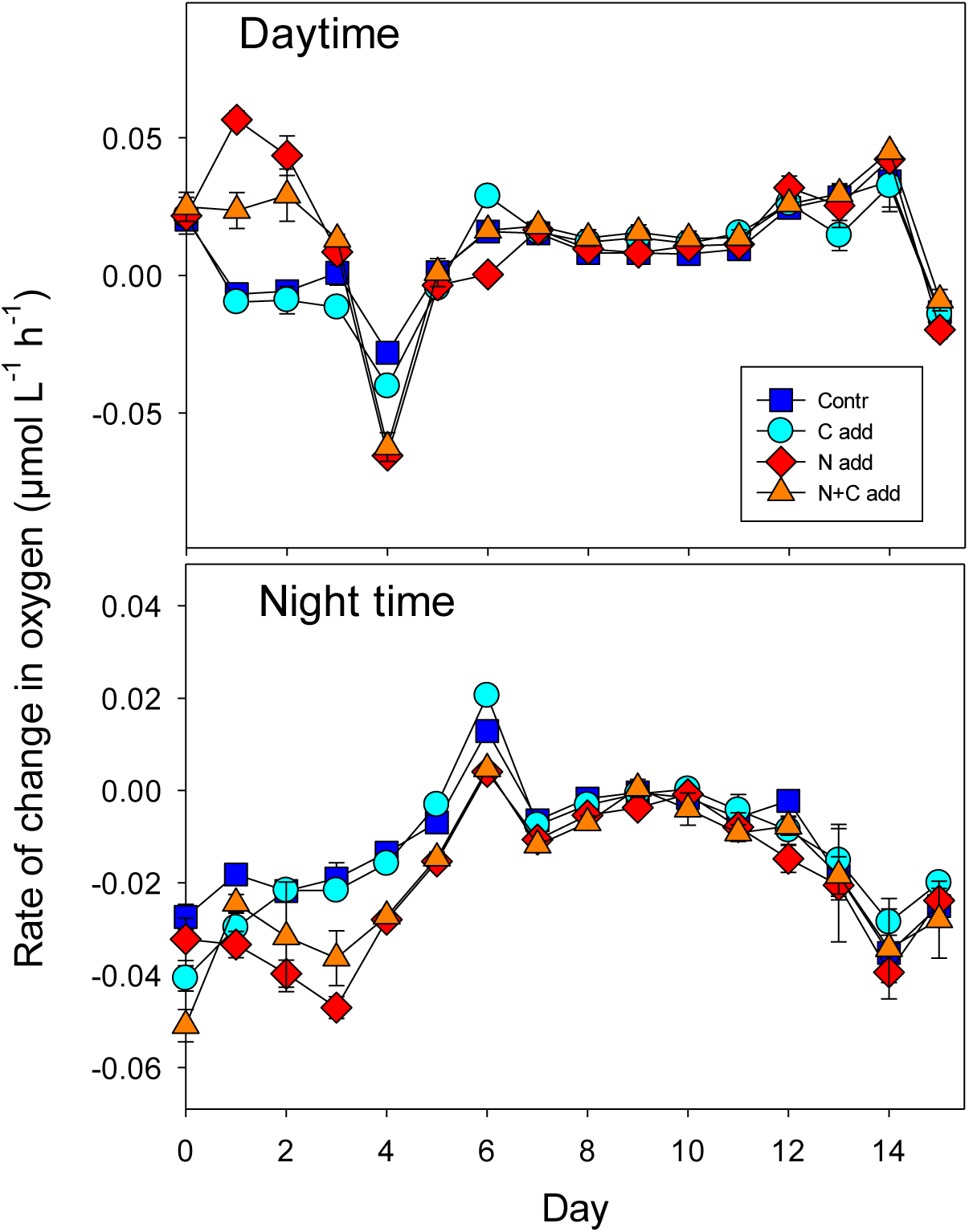
Rate of change in dissolved oxygen during day- and nighttime.

**Supplementary Fig S7.**
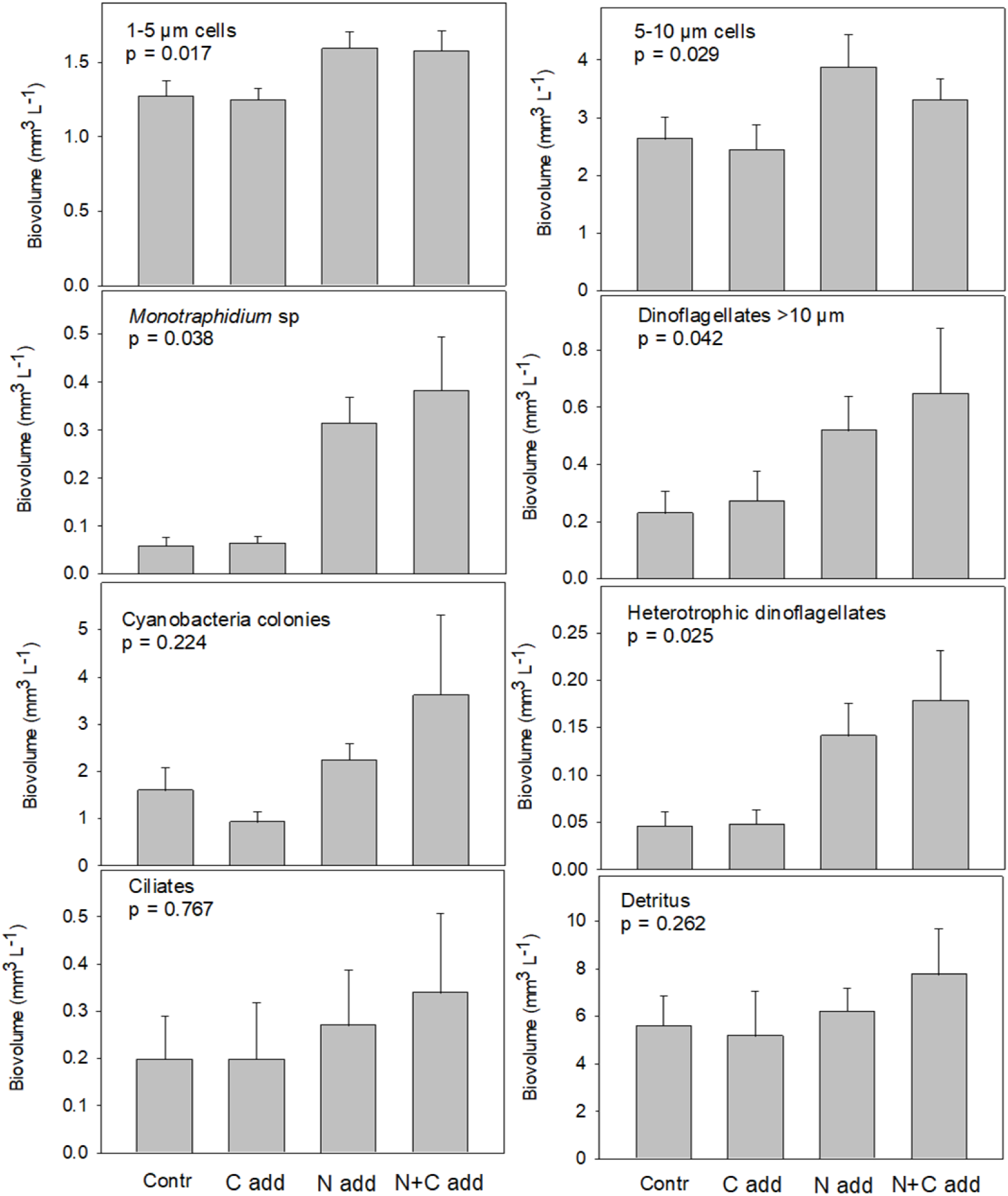
The sum of all biovolume measurements throughout the experiment of different nano- and microplankton groups. The presented p-value is an ANOVA statistical test comparing the different treatments. The temporal development in biovolume is presented in Fig 5.

